# Myelin insulation as a risk factor for axonal degeneration in autoimmune demyelinating disease

**DOI:** 10.1101/2021.11.11.468223

**Authors:** E Schäffner, JM Edgar, M Lehning, J Strauß, M Bosch-Queralt, P Wieghofer, SA Berghoff, M Krueger, M Morawski, T Reinert, W Möbius, A Barrantes-Freer, M Prinz, DS Reich, A Flügel, C. Stadelmann, R Fledrich, RM Stassart, KA Nave

## Abstract

Axonal degeneration determines the clinical outcome of multiple sclerosis (MS), and is thought to result from exposure of denuded axons to immune-mediated damage. We challenge this view after finding in MS and its mouse models that myelin itself increases the risk of axons to degenerate under inflammatory conditions. We propose a model for demyelinating diseases in which for axons that remain myelinated, and thus shielded from the extracellular milieu, dependence from oligodendroglial support turns fatal in an autoimmune disease environment.

---

Demyelination is widely considered the principal cause of axonal degeneration in multiple sclerosis (MS) lesions. However, axonal pathology is an early feature of human MS and experimental autoimmune encephalomyelitis (EAE), with focal axonal damage observable within hours after EAE onset, suggesting that calcium-mediated axon injury precedes overt loss of myelin^1–6^. Despite this, and based on the assumption that de- and amyelinated axons are especially vulnerable to injury, initial damage in MS is proposed to occur at areas naturally devoid of myelin, such as nodes of Ranvier^4–6^. We and others hypothesized that oligodendrocytes provide trophic support to axons^7–9^ because axons are isolated from extracellular glucose at least in part by myelin itself^10^. Consequently, fully myelinated axons should be most dependent on oligodendrocyte– mediated metabolic support, which, in turn, would make myelinated axons most vulnerable to autoimmune attacks that impact the oligodendrocyte’s capacity to support the myelinated axon.

We first sought to reassess the relationship between demyelination and axonal damage in human MS, as only sparse data on the ultrastructural characteristics of axonal pathology in MS lesions are available. To this end, we performed electron microscopy on n = 4 MS patient’s biopsies, which exhibited severe to mild demyelination and inflammation (**Fig.1a,b and Ext.Fig.1a-c, Ext.Fig.2a**). In all lesion types, axonal pathology could be assigned to one of two categories: (I) axonal swellings with intra-axonal organelle accumulations or (II) axons with highly condensed axoplasm (**Fig.1a, Ext.Fig.1a-c, Ext.Fig.2a,b**). Axonal swellings were observed previously in myelin mutant mice and shown to correlate with axonal transport deficits, secondary to a loss of oligodendroglial support of the myelinated axons^11–14^. In addition, early signs of impaired axonal transport were also described in EAE^4,5^. While we found axonal organelle accumulations relatively more frequently in extensively demyelinating lesions, axons with highly condensed cytoplasm showed a reciprocal correlation, being relatively more abundant in lesions with less pronounced demyelination (**Fig.1b,c**). Notably, when assessing the myelination status of the two axonal pathologies, we found all axons with condensed axoplasm to be myelinated (**Fig.1d**) and to be characterized by a thick myelin sheath indicative of preserved myelin rather than remyelination (**Fig.1a and Ext.Fig.1c**). In contrast, organelle accumulations were prevalent in both, myelinated and demyelinated axons (**Fig.1c**). Yet, also many completely demyelinated axons appeared normal (**Fig.1a**).

**Fig. 1:**
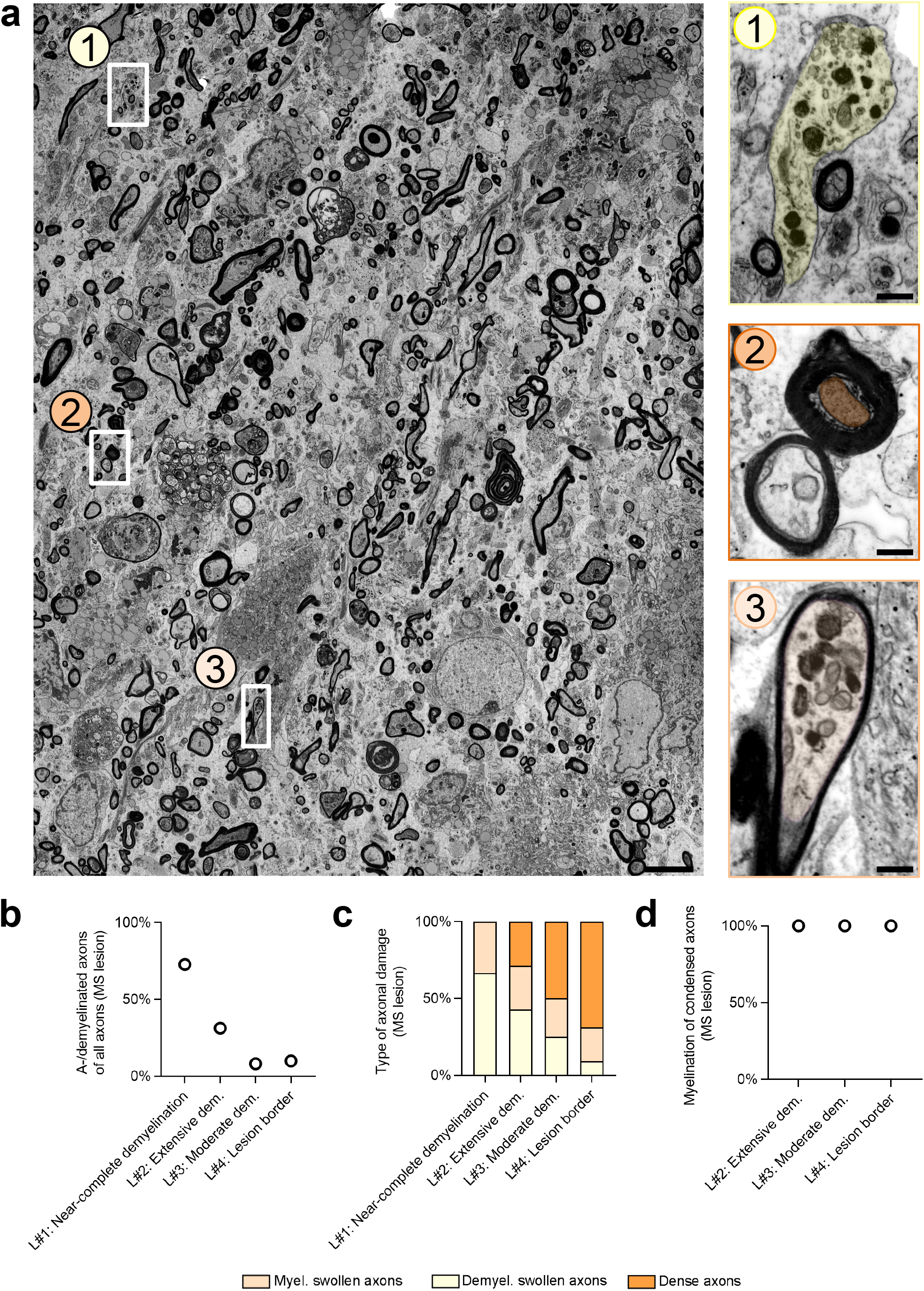
In human multiple sclerosis lesions, irreversible axonal damage prevails in myelinated fibers. **a,** Representative electron micrograph of a human MS lesion with moderate demyelination activity. Common types of axonal damage are exemplified, including demyelinated swollen axons, (highlighted in light yellow, blow-up in 1), axons with condensed axoplasm (dark orange, 2) and myelinated swollen axons (orange, 3). Scale bar of **a**: 5μm and of blow-ups **1-3**: 0.5μm. **b**, Quantification of the extent of demyelination in four individual human MS lesions (L#1 lesion with near-complete demyelination, L#2 active lesion with extensive demyelination, L#3 active lesion with moderate extent of demyelination and L#4 lesion border with limited demyelination). **c,**Quantification of the abovementioned damage types shows an inverse correlation between the extent of demyelination in the 4 MS lesions and the percentage of irreversibly damaged axons. **d,** Quantification of the myelination status of condensed axons reveals that irreversibly damaged axons are always myelinated, independent of the lesion characteristics.

Of note, we furthermore observed possible “transition” types between both axonal pathologies, marked by organelle accumulations and condensed axoplasm, in myelinated axons (**Ext.Fig.2b**).

As condensed axoplasm marks irreversibly damaged axons^15,16^, whereas focal swellings and organelle accumulations may be potentially reversible^4^, we hypothesized that axonal organelle accumulations represent an early consequence of autoimmune-mediated oligodendrocyte dysfunction. To determine the temporal dynamics of axonal pathologies, we induced experimental autoimmune encephalomyelitis (EAE) in C57/BL6 mice by immunization with MOG_35-55_ (**Suppl.Fig.1a-e**). In this variant of EAE, B-cells and antibodies play only a minor role, and demyelination is moderate in the acute disease phase^17,18^. We first analyzed EAE lesions in the ventrolateral lumbar spinal cord at disease peak (4 days post disease onset), when immune cell infiltration is prominent throughout the lesion (**Suppl.Fig.1b-e**), and assessed axonal pathology. At this early disease time point, numerous axons appeared abnormal, and we found by electron microscopy (**Fig.2a,b**) as well as by immunohistochemistry and confocal analysis (**Fig.2c,d**) that pathology was most prevalent in myelinated axons. Axonal loss was unlikely to account for this phenomenon at this early time point, because the total number of axons per area, when corrected for immune cell occupancy, was not reduced in EAE animals compared to controls and the majority of phagocytosed axons harbored residual myelin sheaths (**Suppl.Fig.1e-g**). What characterizes injury of myelinated fibers during the early phase of EAE? In addition to entire myelin-axon profiles engulfed by phagocytosing cells (**Suppl.Fig.1f,g**), we detected the same two types of axonal pathologies as previously: axonal swellings with organelle accumulations and axons with highly condensed axoplasm (**Suppl.Fig.1h**), recapitulating our observations in MS. When we examined the temporal evolution of axonal pathologies in EAE, we found the number of axon cross-sections with organelle accumulations to be highest at very early time points and to subsequently gradually decrease (**Fig.2e**). Conversely, the percentage of axon cross-sections with highly condensed axoplasm increased during the same time course (**Fig.2f**). Likewise, during late stage EAE, almost all condensed axons were myelinated, even at 40 days post EAE induction and in areas of otherwise overt demyelination (**Fig.2g, Suppl.Fig.2a**). In contrast, the myelination status of organelle-filled axonal swellings changed dynamically throughout the disease course decreasing from over 90% at 3 days post disease onset to 40% at 40 days post disease induction (**Fig.2h**). Incidentally, electron microscopy of EAE nerve fibers in longitudinal sections revealed that where organelle accumulations occurred in axonal segments with remaining myelin sheaths, adjacent, demyelinated segments appeared normal (**Ext.Fig.3a**); consistent with our findings in MS biopsies and in toxin-induced models of demyelination (**Ext. Fig.3b-d**).

**Fig. 2:**
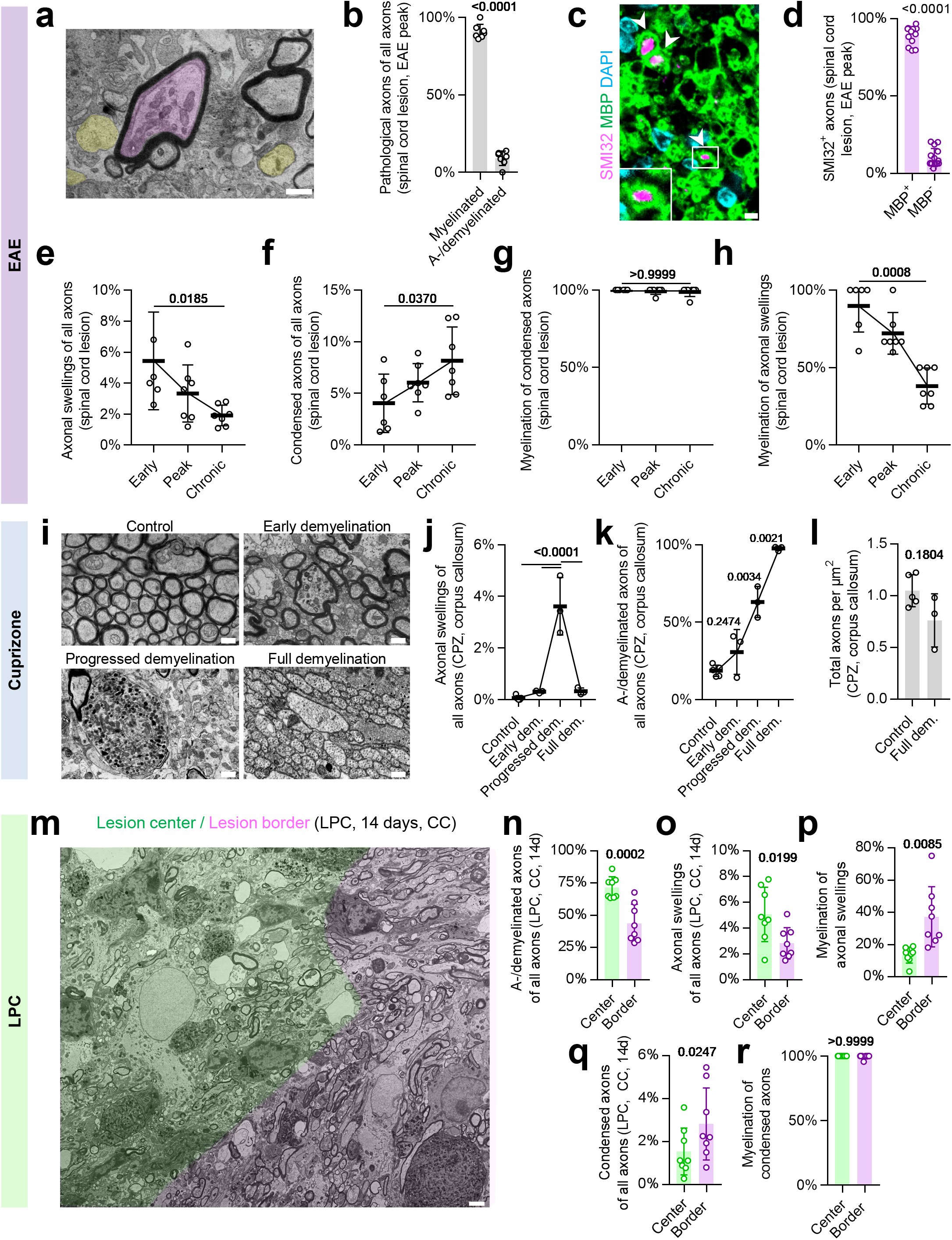
The dynamics of axonal pathology in EAE and toxin-induced models of demyelination. **a,** Representative image of an experimental autoimmune encephalomyelitis (EAE) lesion within the ventrolateral lumbar spinal cord at disease peak (4 days post symptom onset, dpo). Axons with signs of pathology (such as mitochondria accumulation, pink) are often myelinated, while amyelinated axons (yellow) are largely intact. Scale bar = 1μm. **b,** Quantification of the percentage of ultrastructurally damaged axons shows that most damaged axons are myelinated at EAE peak. Unpaired two-tailed Welch’s t test. **c,** Representative images of a spinal cord lesion at 4dpo immunostained against SMI32 (pink), a marker for dephosphorylated neurofilaments, and myelin basic protein (MBP, green). Arrowheads show SMI32^+^ axons with surrounding MBP^+^ myelin sheaths. Scale bar = 2.5μm. **d,** The vast majority of SMI32^+^ axons are myelinated as revealed by quantification of either MBP positive or MBP negative SMI32^+^ axons. Unpaired two-tailed Welch’s t test. **e,f**, Quantification of axon pathology in electron micrographs of ventrolateral lumbar spinal cord lesions 3 dpo (early, n = 6), 4 dpo (peak, n = 7) and 40 days post induction (chronic, n = 7) reveals that while axonal swellings become less frequent over time, condensed axons accumulate throughout the disease course. **g,** Of note, axons with highly condensed axoplasm are almost always myelinated. **h,** In contrast, the percentage of myelinated axonal swellings decreases along with EAE disease progression. **e-h:** One-way-ANOVA with Tukey’s multiple comparisons test (e-f) or Kruskal-Wallis test with Dunn’s multiple comparisons test (g-h) (p-value for early vs. chronic EAE). **i**, Representative images of the corpus callosum without cuprizone treatment (Control, n = 5), cuprizone feeding for 3 weeks (early demyelination, n = 3), for 5 weeks (progressed demyelination, n = 3) and 12 weeks (full demyelination, n = 3). Scale bar = 1.5μm. **j,** While the percentage of axonal swellings first strongly increases along with cuprizone (CPZ) lesion development, only few axonal swellings characterize fully demyelinated CPZ lesions as revealed by quantification of electron micrographs. One-way-ANOVA with Tukey’s multiple comparisons test (p-values shown for comparison with progressed demyelination time point). **k,** Quantification of the percentage of a-/demyelinated axons in CPZ lesions after the abovementioned time periods of cuprizone feeding. One-way-ANOVA with Tukey’s multiple comparisons test (p-values shown for comparison in chronological order). **l,** Quantification of axonal numbers in CPZ lesions shows only a mild, non-significant axonal loss after 12 weeks of cuprizone treatment compared to non-treated controls. Unpaired two-tailed Welch’s t test. **m,** Representative image of a 14-day-old lysolecithin lesion within the corpus callosum. Based on the extent of demyelination, lesions have been divided into efficiently demyelinated lesions centers (green) and incompletely demyelinated lesion borders (pink). Scale bar = 2.5 μm. **n,** Accordingly, the quantification of the extent of demyelination reveals a higher percentage of demyelinated axons in the lesion center compared to the border (n = 8). **o,p** Quantification of the percentage of axonal swellings shows a higher abundance in the lesion center (**o**), while axons with condensed axoplasm are more abundant within the lesion border (**p**). **q,r,** Quantification of the myelination status of axonal pathology reveals that axonal swellings are a feature of both, myelinated and demyelinated fibers (**q**), while condensed axons virtually always have a myelin sheath in both, lesion center and border (**r**). **n-r:** Unpaired two-tailed Welch’s t test (n-p) or Mann-Whitney test (r). Circles in the graphs represent biological replicates. Data are shown as mean ± SD. Numbers represent p-values.

Can complete demyelination improve axonal integrity, resolve organelle accumulations, and allow axons to escape from irreversible injury? To address this, we took advantage of the cuprizone model (CPZ) of toxin-induced callosal demyelination in C57/BL6 mice in which demyelination follows a consistent time course (**Fig.2i,j**). In line with our hypothesis, upon cuprizone feeding, organelle-filled axonal swellings appeared and initially increased but then declined with ongoing demyelination (**Fig.2j,k; Suppl.Fig.2b**). Indeed, fully demyelinated lesions contained only very few residual organelle accumulations (**Fig.2j**), supporting the concept that axonal organelle accumulations constitute an early and potentially reversible axonal injury response. Consistent with this, the total number of axons decreased only slightly across the time course of cuprizone-induced (complete) demyelination (**Fig.2l**). Notably, in these cuprizone lesions we detected a fraction of axons (2-4%) with highly condensed axoplasm, which, analogous to our findings in EAE, was a specific feature of (remaining) myelinated axons throughout the disease course (**Suppl.Fig.2c**). We next hypothesized that if late stage, irreversible axonal damage would indeed be the consequence of myelin retention, this type of axonal pathology should predominate in areas with incomplete demyelination compared to areas of efficient myelin loss. To address this assumption, we assessed axonal pathology in the lysolecithin (LPC) model of focal demyelination which allows comparison of the efficiently demyelinated lesion center to the lesion border with only partial myelin loss (**Fig.2m,n**). While we detected numerous axonal organelle accumulations at both sites (**Fig.2o,p**), degeneration-prone axons with highly condensed axoplasm were relatively more abundant at the lesion border (**Fig.2q**). Importantly, also in the lysolecithin model, the condensed type of axonal pathology was restricted to myelinated fibers (**Fig.2r**).

Collectively, our findings lead us to hypothesize that contrary to general belief, persisting myelin ensheathment, in the context of demyelination, does not provide axonal protection, but conversely, poses a risk for axon survival. Hence, are a- and demyelinated axons relatively protected from axonal damage in an acute autoimmune demyelinating setting? To directly test this, we took advantage of a new mouse mutant with reduced expression of myelin basic protein, termed hypomorphic Mbp, *hMbp* mouse^19^, **Fig.3a, Suppl.3a**).

**Fig. 3:**
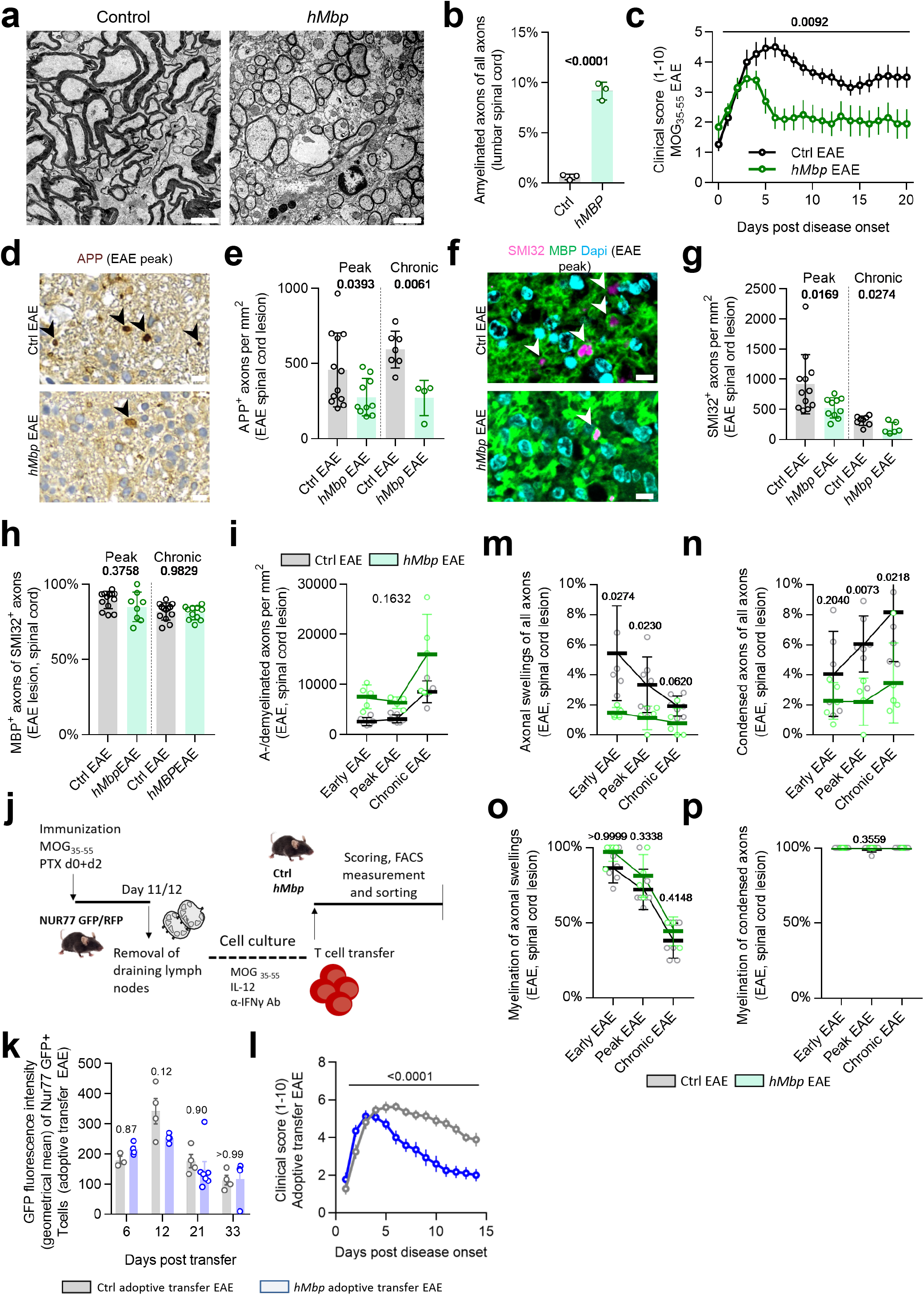
*hMbp* mice demonstrate an amelioration of the clinical disease course and reduced axonal pathology in EAE. **a,** Representative electron micrographs of the ventrolateral spinal cord of control mice and hypomyelinated *hMbp* mice. Scale bar = 2.5 μm. **b,** Quantification of the percentage of amyelinated axons on electron microscopic level reveals around 10% amyelinated fibers in the in lumbar spinal cord in *hMbp* mice. Unpaired two-tailed Welch’s t test. **c,** Clinical EAE scoring (1-10) shows an ameliorated disease course in *hMbp* mice compared to wildtype controls. n Ctrl EAE = 13, n hMbp EAE = 10. Two-way repeated measures ANOVA. **d,** Representative images of APP immunohistochemistry for axonal damage in EAE lesions. Arrowheads point at APP^+^ axons. Scale bar = 10μm. **e,** Quantification of the number of APP^+^ axons in EAE lesions reveals a reduced number of APP^+^ axons in *hMbp* mice at EAE peak (4 days post onset) and chronic EAE (40 days post induction) compared to wildtype EAE controls. Unpaired two-tailed Mann-Whitney test for each time point. **f**, Representative immunohistochemical images of SMI32^+^ MBP^+^ axons in EAE lesions. SMI32^+^ axons are marked by arrowheads and are often surrounded by MBP^+^ myelin sheaths. Scale bar = 5μm. **g,** Quantification of SMI32^+^ axons reveals a reduced number of SMI32^+^ axons in *hMbp* mice at peak and chronic EAE when compared to wildtype EAE controls. Peak EAE: Ctrl n = 12, *hMbp* n = 10, chronic EAE: Ctrl n = 8, *hMbp* n = 6. **h**, The percentage of myelinated SMI32^+^ axons is comparable between *hMbp* and wildtype mice, despite the per se increased number of amyelinated axons in *hMbp* mice. Peak EAE: Ctrl n = 14, *hMbp* n = 8, chronic EAE: Ctrl n = 12, *hMbp* n = 11. Unpaired two-tailed Mann-Whitney test for each time point. **i,** Quantification of a-/demyelinated axons per mm² in electron micrographs of EAE lesions shows that the extent of demyelination between both genotypes is similar. Note that demyelination is most prominent in the chronic EAE phase in both genotypes. Early EAE: Ctrl n = 6, *hMbp* n = 5, peak EAE Ctrl n = 7, *hMbp* n = 5, chronic EAE: Ctrl n = 7, n *hMbp* = 4. Linear regression with slope equality testing. **j,** Schematic outline of the adoptive transfer EAE experiment. **k,** Quantification of the mean fluorescence intensity of GFP (geometric mean) of transferred CD4^+^ T cells as a measure for T cell activation by FACS reveals no difference in T cell activation between *hMbp* mice and controls. Two-way-ANOVA with Sidak’s multiple comparisons test. **l,** Clinical EAE scoring (1-10) after adoptive transfer shows an ameliorated disease course in *hMbp* mice compared to controls, reminiscent of the findings in (**c**). n Ctrl = 14, n *hMbp* = 18. Two-way ANOVA. **m,** Quantification of the percentage of axonal swellings in electron micrographs of EAE lesions shows a reduction in *hMbp* mice compared to EAE. **n**, The respective quantifications of axons with condensed axoplasm demonstrates a lower percentage of condensed axons in *hMbp* mice compared to controls at peak and chronic EAE time points. **o,** Quantification of the myelination status of axonal swellings in *hMbp* and control mice reveals no difference between both genotypes. **q,** In both *hMbp* and wildtype controls, condensed axons are an almost exclusive feature of myelinated axons, independent of the in general increased percentage of amyelinated fibers in *hMbp* mutants. **m-p:** Unpaired two-tailed Welch’s t test for each time point. Circles in the graphs represent biological replicates. Data are shown as mean ± SD, apart from clinical scores (c,k) which are shown as mean ± SEM. Numbers represent p-values.

These mice exhibit an increased fraction of amyelinated axons throughout the CNS, whereas the number and size of all axons appear unaffected (**Fig.3b, Suppl.Fig3b).** Specifically, in the lumbar spinal cord, *hMbp* mice demonstrate around 10% of amyelinated, predominantly medium-sized axons (compared to around 2% in controls),(**Fig.3b, Suppl.Fig.3c,d**). Remarkably, upon EAE induction, *hMbp* mice showed a less severe clinical course of disease compared to respective wildtype controls (**Fig.3c**). Moreover, axonal damage was significantly lower in EAE lesions of *hMbp* mice compared to wildtype controls, at the acute and chronic disease phases, when assessed by immunohistochemistry against APP and de-phospohorylated neurofilament (SMI32) (**Fig.3d-g**). Importantly, we observed no differences in lesion numbers, lesion sizes or localization in *hMbp* mutants compared to controls (**Suppl.Fig.3e-g**). Moreover, the extent of demyelination was similar between both groups and throughout the disease course (**Fig.3i**). To rule out underlying immunization differences, we compared the immunological response of *hMbp* EAE mice to respective wildtype EAE controls using FACS analyses and immunohistochemistry. We found no major differences with respect to T cell and myeloid cell subtypes and numbers at disease peak (**Suppl.Fig.4a-h**). To independently verify these results, we also performed a transfer EAE experiment employing T cells which carry a constitutive red fluorescent marker in addition to an activation-dependent biosensor (Nur77::GFP+::RFP+ reporter mice^20^, **Fig.3j**). While RFP is ubiquitously expressed, T cell re-activation within the CNS tissue triggers a transient GFP expression in transferred T cells, which can be evaluated by flow cytometry. The T cellular infiltration and activation patterns were not significantly different in control vs. *hMbp* mice (**Fig.3k**). Importantly, when following the clinical disease course in *hMbp* and control animals upon adoptive transfer, we confirmed the significantly ameliorated disease course in *hMbp* mutants (**Fig.3l**). This confirms that the improved clinical course and reduced axonal damage cannot be explained by a difference of the immune response in *hMbp* mice. This observation prompted us to also investigate the distribution of axon damage within EAE lesions of *hMbp* mutants. Here, despite the higher number of amyelinated axons, axonal damage remained largely restricted to myelinated axons, as shown by co-immunostaining for SMI32 and MBP (**Fig.3f-h**). Consistent with this, the percentages of axons with organelle accumulations and highly condensed axoplasm were both lower in *hMbp* mice than in controls while the myelination status of axons with either pathology did not differ between genotypes and followed a similar time course (**Fig.3m-p**).

Taken together, our data support a revised working model for inflammatory demyelinating lesions. We hypothesize that the normally symbiotic relationship between the axon and the myelinating oligodendrocyte turns into a fatal one upon disease onset, when the injured oligodendrocyte can no longer provide support of the myelinated axon (**Supp.Fig.5**). Indeed, myelin per se does not support axon integrity^12,13,21^ and, as we demonstrated here, increases the risk of axon degeneration in an autoimmune inflammatory environment. Consistent with previous studies^4,5,22,23,24,^ we found that potentially reversible, locally evolving axonal swellings and transport stasis in not-yet demyelinated axons characterize early diseases stages. Notably, we identified demyelination per se to determine the axon’s fate - by preventing the progression of axonal swellings to irreversible axonal degeneration.

In contrast to the prevailing concept that denuded axons are especially at risk in MS and its models, we here show that irreversible axonal damage is an almost unique feature of myelinated axons in autoimmune lesions, even in experimental models with a basal higher number of amyelinated axons. The concept of axonal pathology resulting from defective axon-glia interactions is in agreement with findings in oligodendroglial mutants^10–13^, and transplantation experiments^14,21,24,25^ and is consistent with our understanding of oligodendroglial-mediated mechanisms, such as axonal metabolic support^7–9,26,27^ and detoxification^28^.

We emphasize that our model of myelin insulation as a risk factor for axonal integrity is not mutually exclusive with recent studies on the role of reactive oxygen/-nitrogen species and intra-axonal calcium levels in mediating axonal injury in EAE^4–6^, as we note that disease mechanisms in autoimmune demyelinating diseases are complex and involve a multitude of intertwining pathomechanistic cascades. Moreover, axons lacking sufficient trophic support by dysfunctional oligodendrocytes are likely to suffer from a reduced capacity to cope with additional injury signals, and thus to show a higher baseline vulnerability compared to their denuded counterparts. Finally, while we suggest rapid and efficient demyelination is protective in early autoimmune disease stages, chronic demyelination may impose independent additional risks to axons and impair axonal function, underlining the importance of promoting remyelination in MS lesions. Taken together, our study suggests that not demyelination per se, but oligodendroglial integrity and downstream axonal support may constitute crucial future therapeutic targets for acute autoimmune demyelinating diseases such as MS.

## Material and Methods

### Human material

Human MS analysis was performed on resin embedded archival brain tissue from human multiple sclerosis patients obtained from the archives of the Institute of Neuropathology, University Medical Center Göttingen, Germany and the Institute of Neuropathology, University Clinic Freiburg. Samples were anonymized and processed in a blinded manner. Epon-embedded tissue was cut into 50 nm sections and processed as described in “electron microscopy”. EM images underwent detailed neuropathological examination and n = 4 lesions of 4 individual patients were selected for further analyses based on the neuropathological criteria of inflammatory infiltrates (active lesions) with either near complete demyelination (L#1), extensive demyelination (L#2) or moderate demyelination (L#3) and a lesion border (L#4) with only minor signs of inflammation. Quantification was performed using FIJI^30^ software. All investigations were performed in compliance with relevant laws and institutional guidelines, and were approved by the ethics committee of the University Medical Center Göttingen and the University Clinic Freiburg.

### Animal models

All mice were bred in a temperature-controlled room with defined 12h light and dark cycles, and food and water *ad libitum*. C57BL6/J mice were used as control animals. *hMbp* mice are described elsewhere in detail and are characterized by >50% reduced expression of MBP^19^. Nur77 GFP+ mice are characterized elsewhere^20^. Mice between 10 and 14 weeks of age were used and all genders were included. For genotyping, DNA was isolated from tail biopsies and incubated in modified Gitschier buffer with Triton-X 100 and proteinase K for 2 hours at 55°C followed by heat inactivation of proteinase K for 10 min at 90°C. Primer sequences are available upon request. All experiments were conducted according either to the Lower Saxony State regulations for animal experimentation in Germany as approved by the Niedersächsische Landesamt für Verbraucherschutz und Lebensmittelsicherheit (LAVES) and in compliance with the guidelines of the Max Planck Institute of Experimental Medicine, Göttingen or approved by the Landesdirektion Sachsen and in compliance with the guidelines of the Paul-Flechsig-Institute, Leipzig.

### Electron microscopy

Mice were perfused with PBS and two lumbar spinal cord segments were dissected and post-fixed in 0.1M phosphate buffer containing 2.5% glutaraldehyde and 4% paraformaldehyde for at least 1 week at 4°C. Pieces were then embedded into Agar 100 and cured for 48 h at 60°C. 500nm semithin sections were stained with a methylene blue / azure II solution for 1 minute on a 60°C heating plate. 50 nm ultrathin sections were taken up with Formvar-coated grids and contrasted with Uranlyess (Delta Microscopies) and lead citrate. Images were taken with a SIGMA electron microscope (Zeiss) equipped with a STEM detector and ATLAS software. For each lesion, at least 15 images of 4000x magnification were analyzed using FIJI^29^ software.

### Classical experimental autoimmune encephalomyelitis

EAE induction and staging were performed as described elsewhere^30^. Mice were kept in the room of the procedure for at least 2 weeks to allow for acclimatization. Briefly, at the day of immunization, mice were anesthetized with isoflurane and an emulsion of Complete Freund’s Adjuvant (CFA) made from Mycobacterium tuberculosis (BD Bioscience, 231141) with a total 200μg MOG_35-55_ (AnaSpec, AS-60130-1) per mouse was injected subcutaneously into all 4 flanks. 400 ng pertussis toxin (Sigma, P7208) was administered intraperitoneally after immunization and 48 hours later. Mice were weighed and scored every day at the same time. Mice that had a disease onset after day 20 were excluded. Mice that exceeded a score of 4 were put on special bedding for easier movement and water bottles and food were put inside the cage for easier accessibility.

### Adoptive transfer EAE

For EAE induction through adoptive transfer of pathogenic T cells, donor Nur77 GFP+ RFP+ mice were immunized with 75 μg MOG_35-55_ per mouse in CFA. 200ng Pertussis toxin was administered intraperitoneally on d0 and d2 post immunization. Draining lymph nodes were harvested on d12 post immunization. A single cell suspension was prepared and the cells were cultivated for 3 days in the presence of 25 μg/ml MOG_35-55_, 25 ng/ml recombinant mouse IL-12 (R&D, 419-ML) and 20 μg/ml α-IFNу-Ab (BioXcell, clone XMG1.2, BE0055). To induce EAE, 3*10^6^ cells were injected intraperitoneally into recipient *hMbp* or wildtype animals with the same genetic background. Clinical signs and body weight were examined daily starting on d4 post T cell transfer. At day 6, 12, 21 and 33 post induction, 3-6 animals were sacrificed and T cell activation was quantified using flow cytometry. Therefore, from n = 14 Ctrl animals undergoing adoptive transfer EAE and clinical scoring, 2 animals were sacrificed at day 7 post disease onset (dpo), 2 animals at day 8 and 1 animal at day 13, hence 9 animals were followed until day 14 dpo. For *hMbp* mutants, a total number of n = 18 animals underwent EAE adoptive transfer, with 3 animals being sacrificed at day 7 dpo and 1 at day 8, hence 14 animals were followed until day 14 post EAE disease onset.

### Immunohistochemistry and imaging

For immunohistochemistry, mice were perfused with PBS and 2 lumbar spinal cord segments were dissected and post-fixed in 4% PFA in PBS overnight at 4°C. Samples were embedded in paraffin and cut into 5μm sections. Samples were de-waxed in xylol and ethanol, antigen retrieval was performed using boiling citrate buffer (pH 6), and samples were blocked with the serum of the secondary antibody. Primary antibodies were applied overnight at 4°C and included SMI31 (Biolegend, 801601), IBA1 (Wako, 019-19741), SMI32 (Biolegend 801701), MBP (Invitrogen PA1-10008), Synaptophysin (Synaptic Systems 101 203), APP (Merck MAB348) and CD3 (Abcam, ab5690). For fluorescent stainings, secondary antibodies were applied for 1 hour (all antibodies were purchased from Dianova: Cy2 goat anti mouse (115-225-071), Cy2 goat anti rat (112-225-143), Cy2 goat anti rabbit (111-225-144), Cy3 goat anti mouse (115-165-071), Cy3 goat anti rat (112-165-003), Cy3 goat anti rabbit (111-165-144), Cy3 goat anti chicken (303-165-003)), counterstained with DAPI (Invitrogen, D1306) and embedded in AquaPolymount (Polysciences, 18606). For chromogenic stainings, samples were labelled with biotin-coupled antibodies (all antibodies were purchased from Southern Biotech: goat anti mouse (1012-08), goat anti rat (3052-08)) for 1 hour at room temperature and labelled structures were visualized using the ABC Vectastain kit (Vector laboratories, PK-6100). Finally, samples were counterstained with hemalum and embedded in Eukitt (O-Kindler). Overview pictures for quantification of bigger structures (e.g. cells) were taken with the Zeiss AxioScan Z1. High-resolution images for more detailed analyses (e.g. SMI32 MBP correlation) were taken with the Zeiss LSM 880 Airyscan confocal microscope. In both cases, images were acquired with Zen imaging software (Zeiss) and analyzed using FIJI^29^.

### Cuprizone-induced demyelination

Cuprizone treatment was performed as described elsewhere^31^. 0.2% w/w Cuprizone (Sigma, C9012) was fed as powder chow and mice were sacrificed at 4 time points: d0 (control without cuprizone), 3 weeks (demyelination onset), 5 weeks (progressed demyelination) and 12 weeks (full demyelination). Food intake and weight were monitored throughout the experiment. At the day of sacrifice, animals were perfused with PBS, followed by 2.5% glutaraldehyde and 4% paraformaldehyde in 0.1M PB for electron microscopy. Fixed brains were cut using a vibratome (Leica VT1200, 300 mm) and the corpus callosum with adjacent tissue was punched with a 2mm punching tool and processed as described for electron microscopy. For electron microscopic analysis and axonal counting, axons smaller than 400nm in diameter were excluded from quantification.

### Lysolecithin-induced demyelination

Stereotactic injection of lysolecithin (Sigma, L4129) was performed as described elsewhere^32^. Mice were anesthetized intraperitoneally with MMF (0.5 mg medetomidin/kg, 5.0 mg midazolam/kg and 0.05 mg fentanyl/kg). The head fur was trimmed and the eyes covered with Bepanthene cream (Bayer). A small incision was made to expose the skull and a hole was drilled at X ± 1.0 mm / Y −0.1 mm from bregma. A capillary filled with 1% lysolecithin and 0.03% Monastral Blue (Sigma, 274011) for lesion visualization was inserted into the brain parenchyma (Z −1.4mm) using a micromanipulator (Nanoliter 2000, World Precision Instruments). 1 μl was injected with a flow rate of 100 nl/min. After the injection, the head skin was sutured and 0.05 mg/kg buprenorphine was administered. Anesthesia was antagonized with AFN (2.5 mg/kg atipamezol, 1.2 mg/kg naloxone, 0.5 mg/kg flumazenil) and mice were closely monitored after the surgery.

### Fluorescent-activated cell sorting standard EAE

Mice were perfused at EAE peak (4 days post onset) with PBS and heparin after blood was taken from the right ventricle into an EDTA pre-filled tube on ice. The spinal cord was dissected and dissociated with a scalpel on a drop of medium containing Hanks’ balance salt solution (HBSS), glucose and HEPES. The cell solution was filtered through a 40 μm strainer, centrifuged, re-suspended in 75% Percoll (GE Healthcare) and layered under a 25% Percoll solution topped with PBS. After centrifugation for 30 minutes with slow breaking, distinct gradients became visible and myelin debris was carefully removed. The remaining cell phase was taken up, cleaned by centrifugation with PBS and stained with viability dye (eFluor 506 Fixable Viability Dye, Thermo Fisher) for 30 minutes on ice. In parallel, erythrocyte lysis (BD Pharma) was performed for blood samples. For both sample types, cells were blocked with Fc block (CD16/32, Invitrogen, 14-0161-82) for 20 minutes on ice. After washing, antibodies were incubated for 20 minutes on ice. For lymphocytes, CD45^+^ CD11b^−^cells were gated and differentiated using CD4 and CD8. CD45^+^ CD11b^+^ myeloid cells were further differentiated into neutrophils (Ly6C^hi^, CD115^−^) and two populations of monocytes (Ly6C^hi^ or Ly6C^lo^, CD115^+^). Following antibodies were used: CD11b-BV421 (BioLegend 101236), Ly6C-Alexa488 (BioLegend 128022), CD4-PE (BioLegend 100512), CD8b-Alexa647 (BioLegend 126611), CD45-APC-e780 (Invitrogen, 47-0451-82), CD115-PE-Cy7 (Invitrogen, 25-1152-82). FACS analysis was performed using a FACS Aria III system (BD Biosciences). Data were analyzed using FlowJo software (Tree Star).

### Fluorescent-activated cell sorting adoptive transfer EAE

For analysis of adoptive transfer EAE spinal cord tissue, mice were perfused as described above and the spinal cord digested with Liberase (Roche) and DNAse I (Sigma) for 15 min at 37°C in the water bath. Subsequently, a single cell suspension was prepared and myelin was removed by Percoll gradient (40%) centrifugation. After centrifugation, the cell pellet was re-suspended in FACS buffer and filtered through a 40 μM filter. The cells were blocked using an Fc blocking antibody (Biolegend Fc-block clone: 2.4G2) and lymphocytes were stained with CD3 and CD4. All FACS antibodies were purchased from Biolegend: CD4 Pe-Cy5 (clone: GK1.5) and CD3e APC (clone:145-2C11). CD4+ T cells (CD3+CD4+) were divided in transferred (RFP+) and endogenous (RFP-) T cells. T cell activation was measured by the level of GFP expression (geom. mean fluorescent intensity) of transferred T cells. The data were analyzed using the FlowJo software (Tree Star).

### Statistical analysis

For Power analysis the software G*Power Version 3.1.7. was used. Power analyses were performed before conducting *in vivo* experiments (a priori). Adequate Power (1 – beta-error) was defined as ≥ 80% and the alpha error as 5%. Data are expressed as individual biological samples with mean ± standard deviation (SD) unless indicated otherwise. All data were processed and statistically analyzed using MS Excel and GraphPad Prism v7.05, unless indicated otherwise. The respective statistical test is indicated in the figure legends. Briefly, normal distribution was tested using the D’Agostino-Pearson omnibus normality test or Shapiro-Wilk test. For comparing two groups, the unpaired two-tailed Welch T-test (normal distribution) or Mann-Whitney test (no normal distribution) was used. For comparing more than two groups, a one-way ANOVA with the Tukey’s multiple comparisons test (normal distribution) or the Kruskal-Wallis test with Dunn’s multiple comparisons test (no normal distribution) were used, and for comparing two or more groups for more than one time point (longitudinal analysis), a two-way ANOVA with the appropriate post-test was used. Statistical differences were considered to be significant when p < 0.05 and indicated as exact numbers in the graphs.

## Data Availability

All relevant data of the present manuscript are available from the corresponding authors on reasonable request.

## Acknowledgements

We are grateful to C. Maack, T. Ruhwedel, A. Fahrenholz (MPI Göttingen), A. Wohltmann (Institute of Neuropathology, Göttingen), T. Rudolf, N. Schwagarus, C. Paul (Paul-Flechsig-Institute Leipzig) and J. Craatz (Institute of Anatomy, Leipzig) for skillful technical support. We also thank T. Pawelz (MPI Göttingen) and J. Meißner, E. Jung, S. Herrmann and P. Hirrlinger (MEZ Leipzig) for excellent animal husbandry. J.Strauss and A.Flügel are supported by DFG grant FL 377/4-1, and A.Flügel is supported by an ERC Advanced Grant (101021345, T-Neuron). D.S.Reich is supported by the Intramural Research Program of NINDS and by the AMRF. E. Schäffner was supported with a Boehringer Ingelheim Foundation travel grant. R.Fledrich is supported by a DFG Emmy-Noether fellowship. R.M.Stassart is supported by an ERC Starting Grant (948857, AxoMyoGlia). K.A.Nave is supported by an ERC Advanced Grant and the AMRF. J. M. E is supported by Grant 127 from the UK MS Society.

## Author contributions

K.A.N. and R.M.S. designed and supervised the study and wrote the manuscript. E.S. performed and planned the experiments and contributed to the manuscript. R.F. contributed to study supervision, manuscript writing and discussions. J.M.E. contributed to study design, discussions and manuscript writing. S.B. and M.L. contributed to cuprizone experiments. M.B.Q. and M.L. contributed to lysolecithin experiments. P.W. contributed to FACS experiments, A.F. and J.S. performed adoptive EAE experiments. M.W. contributed hMbp mice. M.M. contributed to immunofluorescence imaging. M.K. and T.R. contributed to electron microscopy experiments, C.S.-N. and M.P. provided human multiple sclerosis samples. A.B.F. and D.R. contributed to discussions and critical review of the manuscript.

## Competing interests

The authors declare no competing interests.

---

**Extended Fig. 1:**
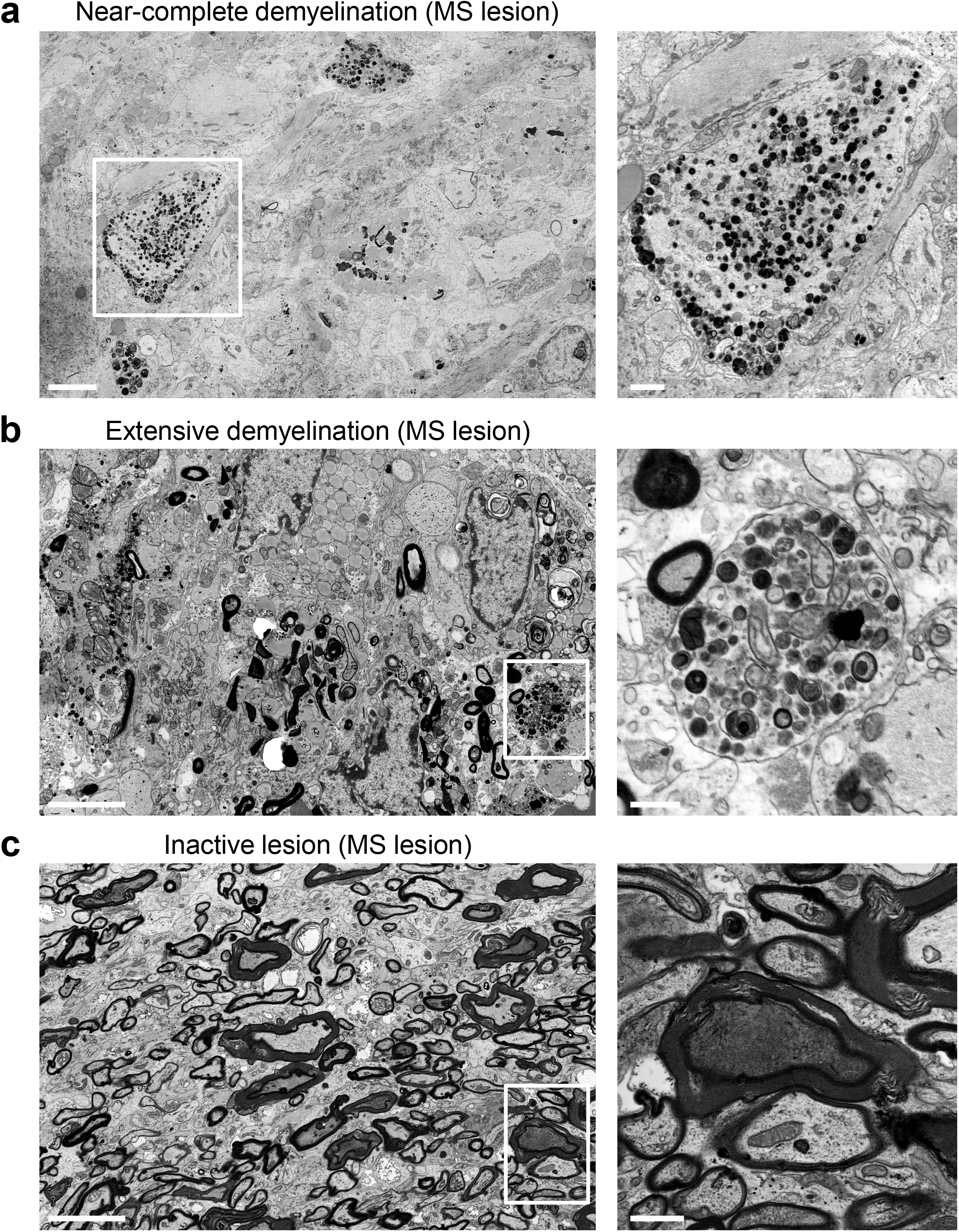
Human multiple sclerosis lesions. **a,** Representative electron micrograph of a lesion with near-complete demyelination. Note the virtual absence of myelinated axons. A swollen demyelinated axon is shown in higher magnification. Scale bar = 5μm / blow-up = 1μm. **b,** A representative image from an active lesion with extensive demyelination demonstrates a few myelinated axons, ongoing phagocytosis and many demyelinated axons. A swollen demyelinated axon is shown at higher magnification. Scale bar = 5μm / blow-up = 1μm. **c,** A representative image of a lesion border with a large proportion of myelinated axons. Note the myelinated axons with a pronounced condensation of the axoplasm, representing irreversible axon damage (high magnification image). Scale bar = 5μm / blow-up = 1μm.

**Extended Fig. 2:**
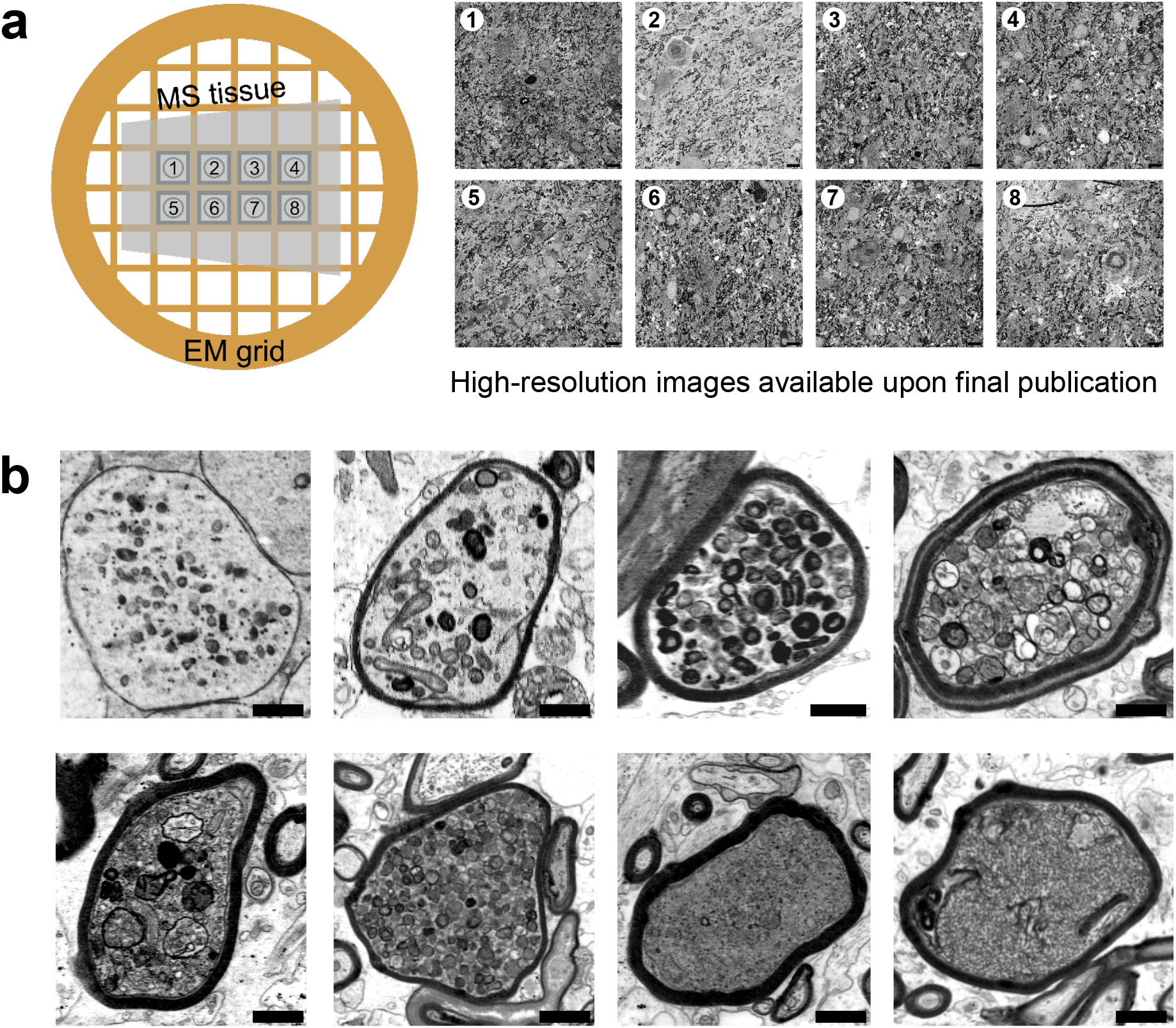
Axonal pathology in human multiple sclerosis. **a,** Eight large, high-quality electron micrographs of an almost complete sample grid of an MS lesion with moderate demyelination showing typical MS lesion characteristics with immune cell infiltration, phagocytosing cells, myelinated and demyelinated axons as well as the different forms of axonal pathology. The full-resolution images are available as a picture set upon final publication. Scale bar = 10μm. **b,** Electron microscopic images of axonal pathology demonstrating the heterogeneity of axonal swellings and axons with highly condensed axoplasm, as well as potential transition types between both forms of axonal damage. Scale bar = 1 μm.

**Extended Fig. 3:**
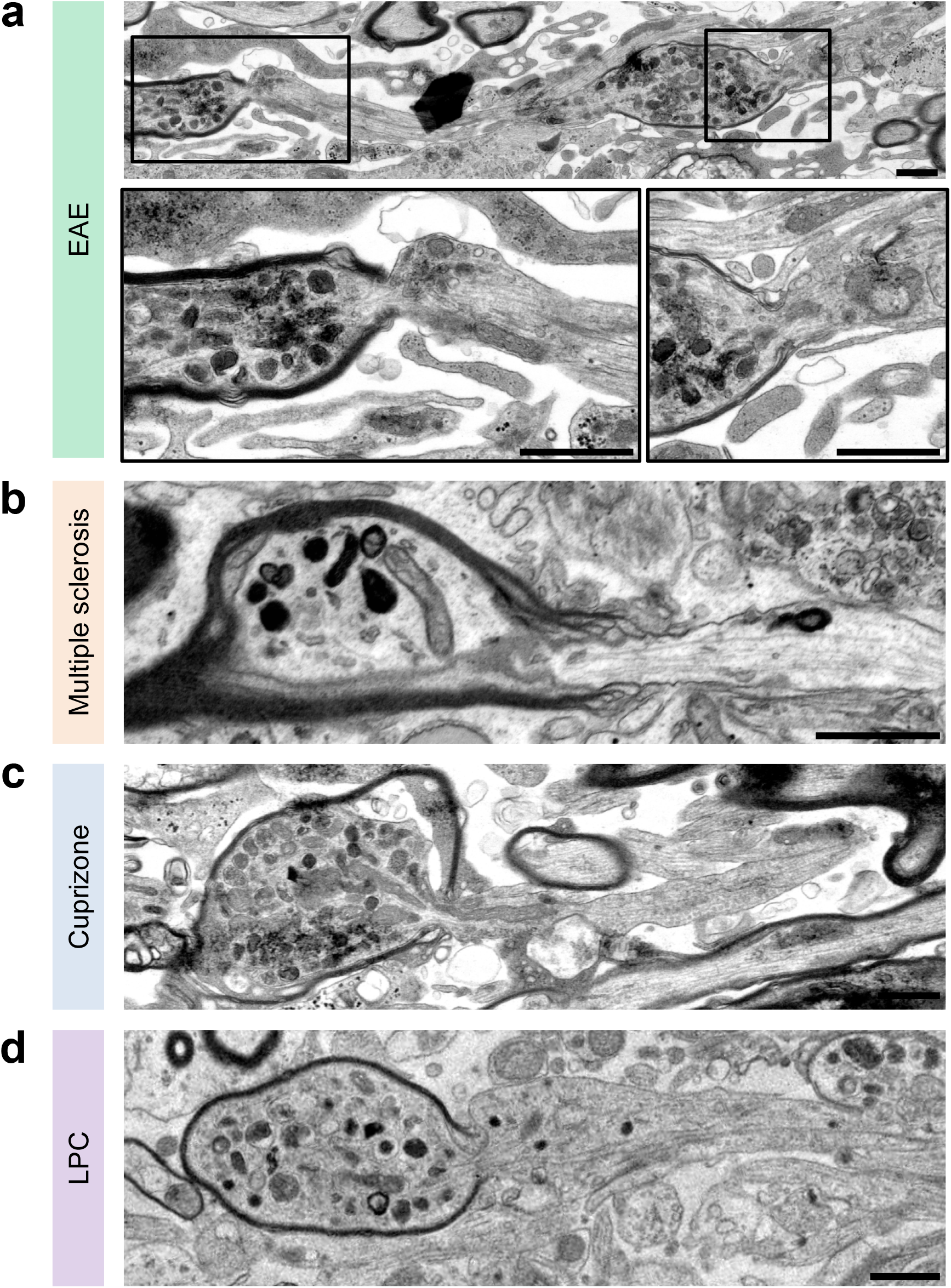
Axonal swellings predominate in myelinated segments. Images of partially demyelinated axons show that organelles accumulate specifically in not or not yet demyelinated parts of the axons. Similar observations were made in **a**, EAE; **b**, multiple sclerosis; **c**, cuprizone and **d**, lysolecithin-induced demyelination. Scale bar = 3μm.

**Suppl. Fig. 1:**
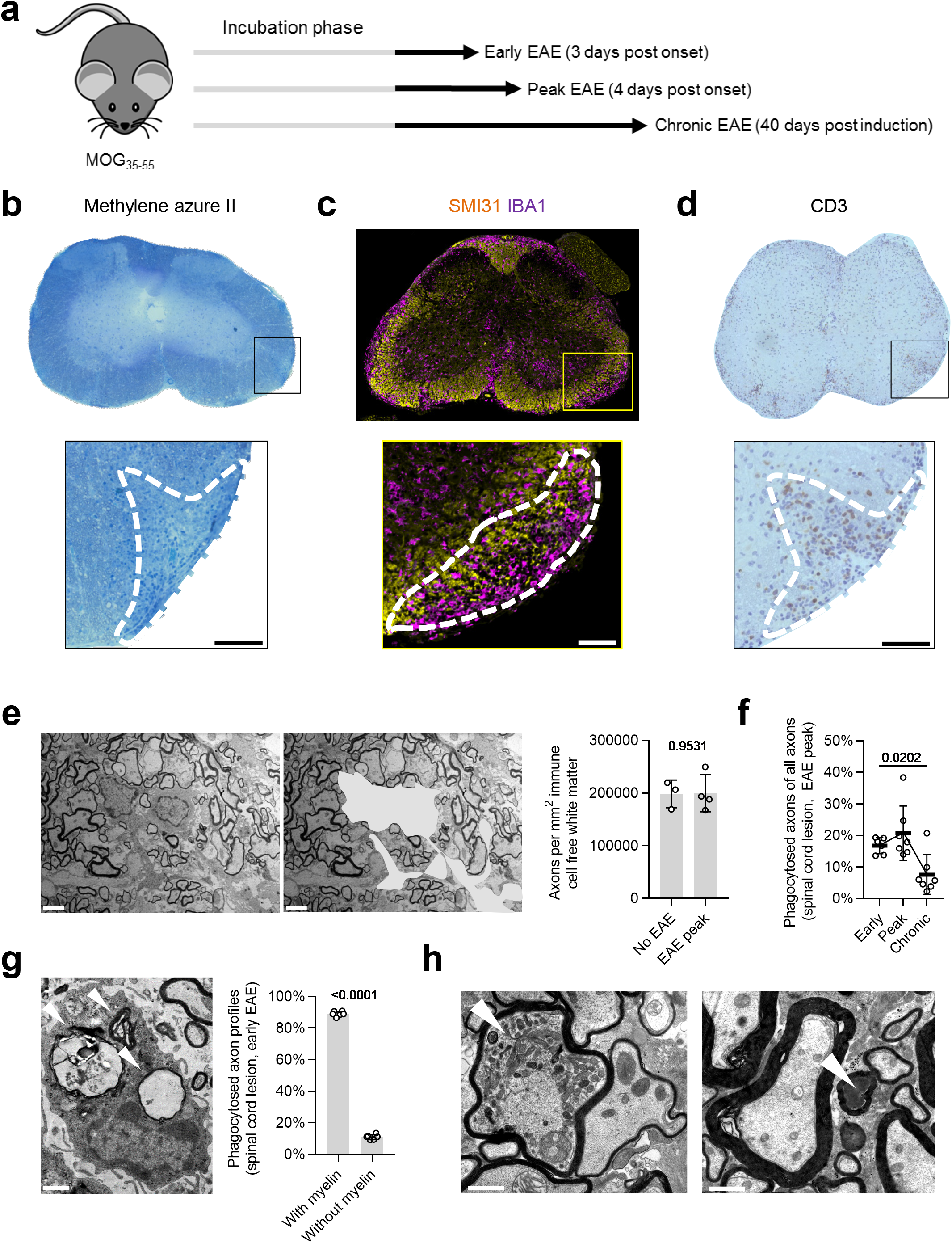
EAE outline and histopathology. **a,** Experimental paradigm of MOG_35-55_ EAE experiments. **b,** Spinal cord lesions are identified in methylene azure II stained semithin sections by accumulation of immune cells in the white matter. The dashed line encircles a lesion. Scale bar = 200μm. **c,** Lesion formation can also be visualized by immunohistochemistry for IBA1, a macrophage/microglia marker and SMI31, a marker for phosphorylated neurofilament. Note the accumulation of IBA1^+^ cells within the lesion (encircled with a dashed line). Scale bar = 200μm. **d,** Representative image of a CD3^+^ staining for T lymphocytes. The dashed line displays a lesion loaded with CD3+ T cells. Scale bar = 200μm. **e,** To estimate axonal numbers in EAE lesions compared to non-EAE controls, the area covered by immune cells was subtracted (right image, grey area) to correct for differences in axonal density due to immune cell occupancy. The respective quantification of axons per mm^2^ shows no significant axonal loss at EAE peak (4dpo). Scale bar = 2.5μm. **f,** Quantification of the percentage of phagocytosed axons reveals that phagocytosis is most pronounced at EAE peak compared to chronic EAE (40dpo). Kruskal-Wallis test with Dunn’s multiple comparisons test (p-value for early vs. chronic EAE). **g**, Representative image of a phagocytosing cell at early EAE. Engulfed axon/myelin profiles are indicated by white arrowheads. Quantification of the myelination state of phagocytosed axons at early EAE reveals that most phagocytosed axons are myelinated. Unpaired two-tailed Welch’s t test. Scale bar = 3μm. **h,** Representative electron micrographs of axonal swellings with organelle accumulations (left image, arrowhead) and irreversible damaged axons with highly condensed axoplasm (right image, arrowhead) in EAE lesions. Scale bar = 3 μm. Circles in the graphs represent biological replicates. Data are shown as mean ± SD. Numbers represent p-values.

**Suppl. Fig. 2:**
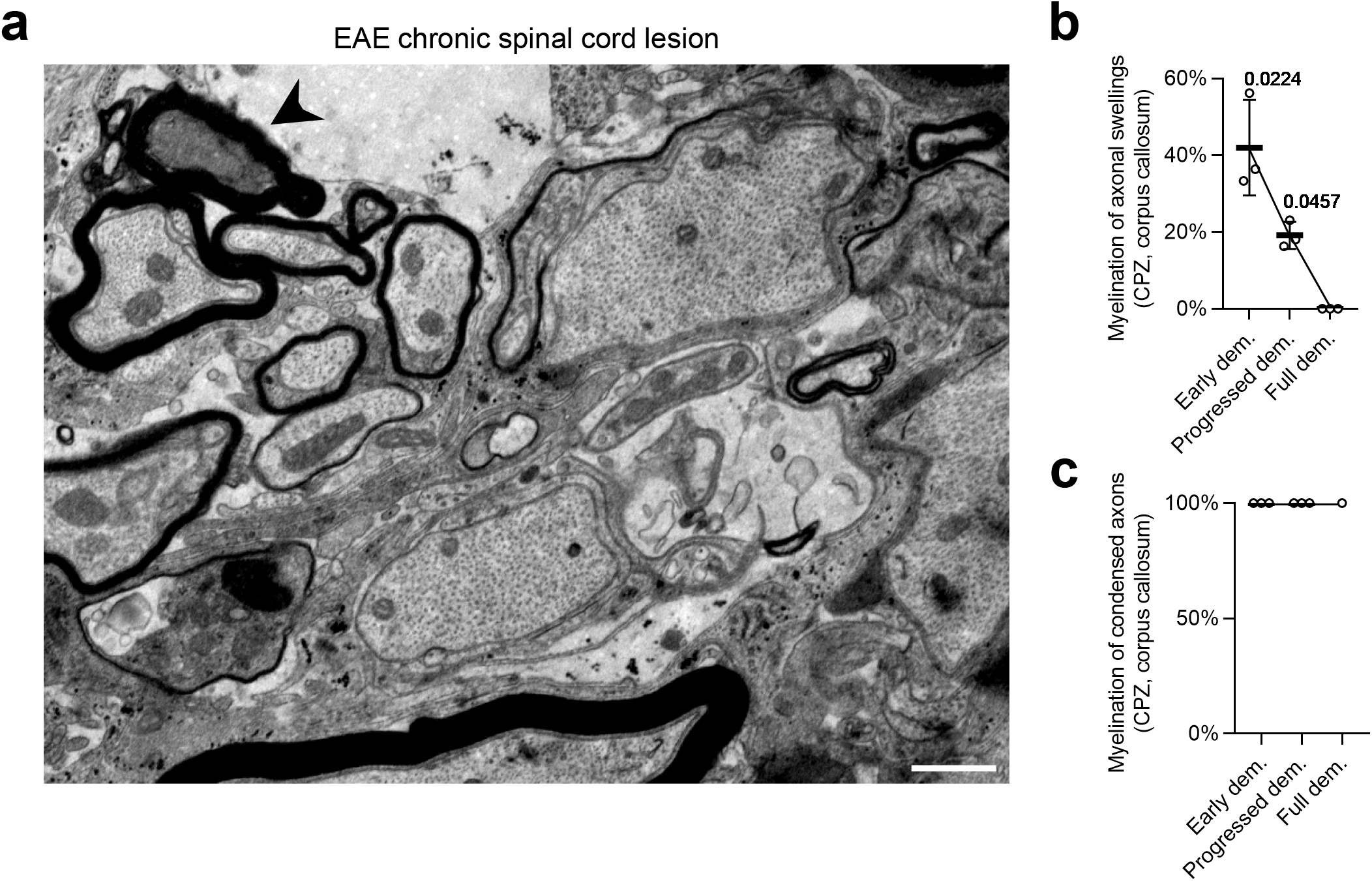
Condensed axons in EAE and the cuprizone model. **a**, A representative electron micrograph of a chronic spinal cord EAE lesion shows that demyelinated axons appear relatively intact, while a proportion of myelinated axons show signs of damage like condensation of axoplasm (arrowhead). Scale bar = 1μm. **b**, Quantification of the myelination status of axonal swellings during cuprizone treatment. Swollen axons become demyelinated over time. One-way-ANOVA with Tukey’s post-test **c**, Quantification of the myelination status of axons with condensed axoplasm during cuprizone treatment. All condensed axons are myelinated. Note that for the full demyelination time point, condensed axons are virtually absent and were only detected in one mouse (n = 1). Circles in the graphs represent biological replicates.

**Suppl. Fig. 3:**
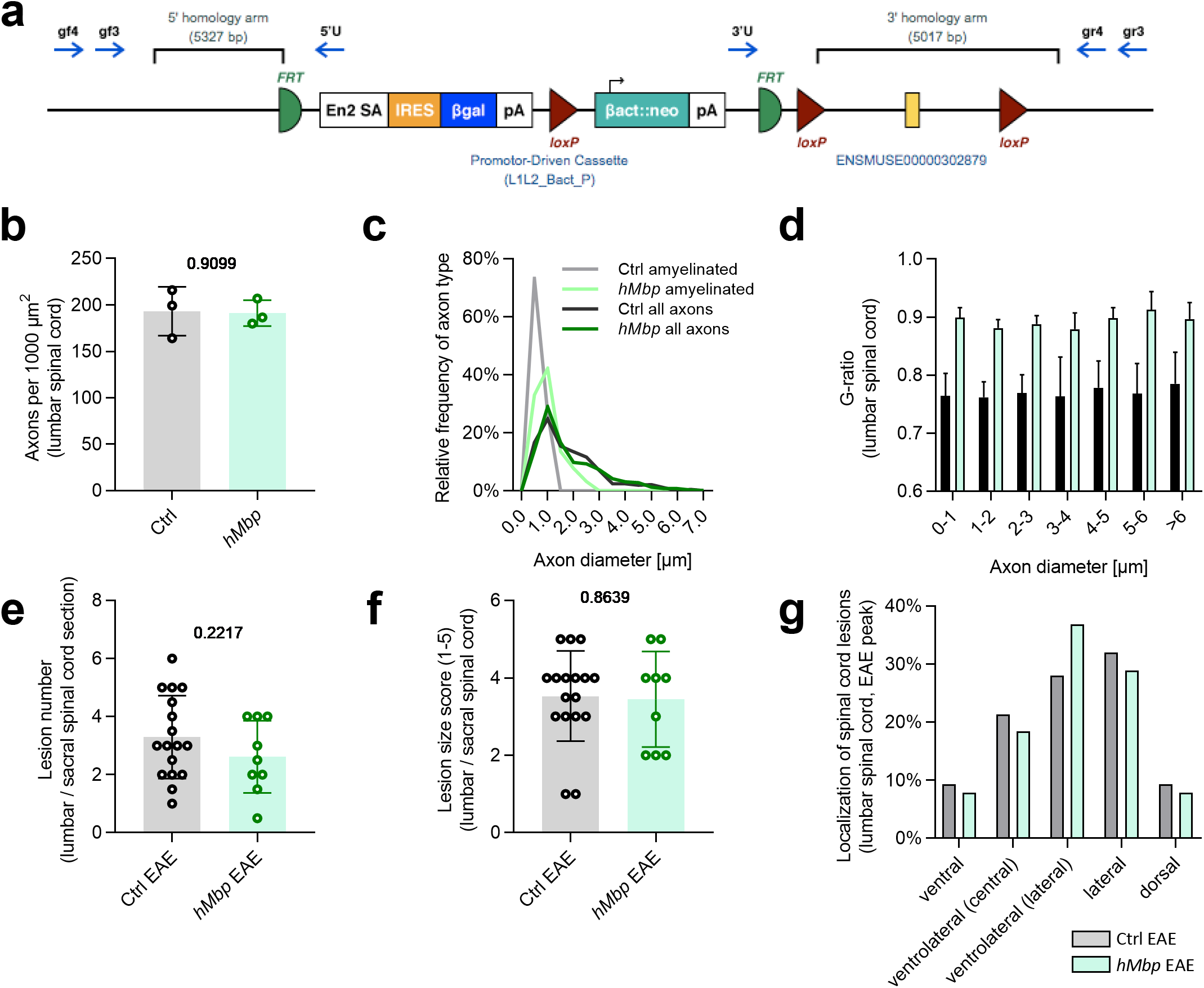
Characterization of hMbp mice. **a**, The genetic construct used in hMbp animals leads to a reduced expression of MBP without the necessity to cross-breed with a Cre-line. **b**, Quantification of axon numbers in the lumbar spinal cord reveals no difference between control and hMbp mice. Unpaired two-tailed Welch’s t test. **c**, Frequency distribution of axonal diameters within the lumbar spinal cord demonstrates similar axon sizes between hMbp and controls, and amyelination of medium-sized axons in hMbp mice that would normally be myelinated. n = 3 in both groups. **d**, G-ratio analysis shows that the hypomyelination of axons in hMbp mice is a feature of all axon calibers. n = 3 in both groups. **e,f**, Spinal cord EAE lesions occur in similar number and frequency in hMbp mice and controls. For each animal, two lumbar sections have been analyzed and the mean is plotted in the diagram. **g**, Lesion localization is comparable in both genotypes. In total, 75 lesions of 17 wildtype animals (mean lesions per animal = 4,4) and 38 lesions of 8 hMbp animals (mean lesions per animal = 4,2) lesions were quantified. Total lesion numbers are higher than in f as lesions can expand to more than one localization. The bars represent the mean distribution of all pooled lesions. In all diagrams, circles in the graphs represent biological replicates. Data are shown as mean ±SD. Numbers represent p-values.

**Suppl. Fig. 4:**
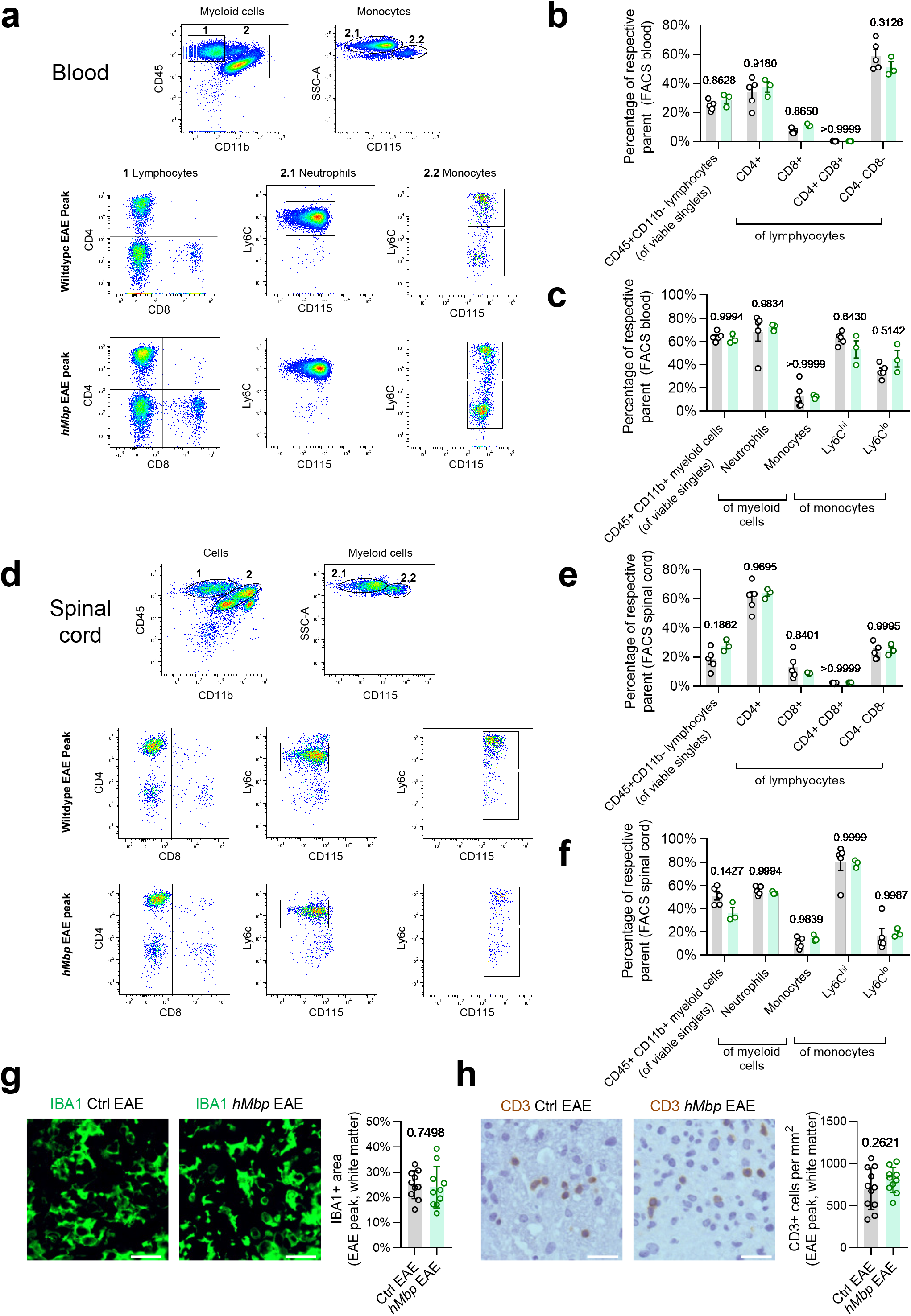
EAE immune cell diversity is similar in control and *hMbp* mice. **a,** Representative flow cytometry of blood samples at EAE peak (4 dpo). Gate 1 = CD45^+^CD11b^−^ lymphocytes, gate 2 = CD45^+^CD11b^+^ myeloid cells (2.1 = CD45^+^CD11b^+^SSc^hi^CD115^−^ neutrophils, 2.2 = CD45^+^CD11b^+^SSc^lo^CD115^+^ monocytes). Ctrl EAE n = 5, *hMbp* n = 3. **b,** Quantification of CD45^+^CD11b^−^CD4^+^ and CD45^+^CD11b^−^CD8^+^ blood lymphocytes reveals no differences in frequencies between *hMbp* mice and wildtype controls. **c**, The distribution of CD45^+^CD11b^+^ myeloid cells based on CD115 and Ly6C expression in blood is similar between both genotypes. **d,** Representative gating of FACS spinal cord cells. Gate 1 = CD45^+^CD11b^−^ lymphocytes, gate 2 = CD45^+^CD11b^+^ myeloid cells (2.1 = CD45^+^CD11b^+^SSc^hi^CD115^−^ neutrophils, 2.2 = CD45^+^CD11b^+^SSc^lo^CD115^+^ monocytes). **e,f** Quantification of lymphocytes and myeloid cell populations in spinal cord reveals no differences in the frequencies between *hMbp* mice and respective controls. **g,** Representative images of IBA1^+^ cells in the lumbar white matter at EAE peak show no differences in IBA1 density, as also revealed by quantification per area. **h**, Representative images of CD3^+^ cells in the lumbar white matter at EAE peak. The quantification shows similar numbers of CD3^+^ cells in the lumbar white matter. Circles in the graphs represent biological replicates. Data are shown as mean ±SD. Numbers represent p-values.

**Suppl. Fig. 5:**
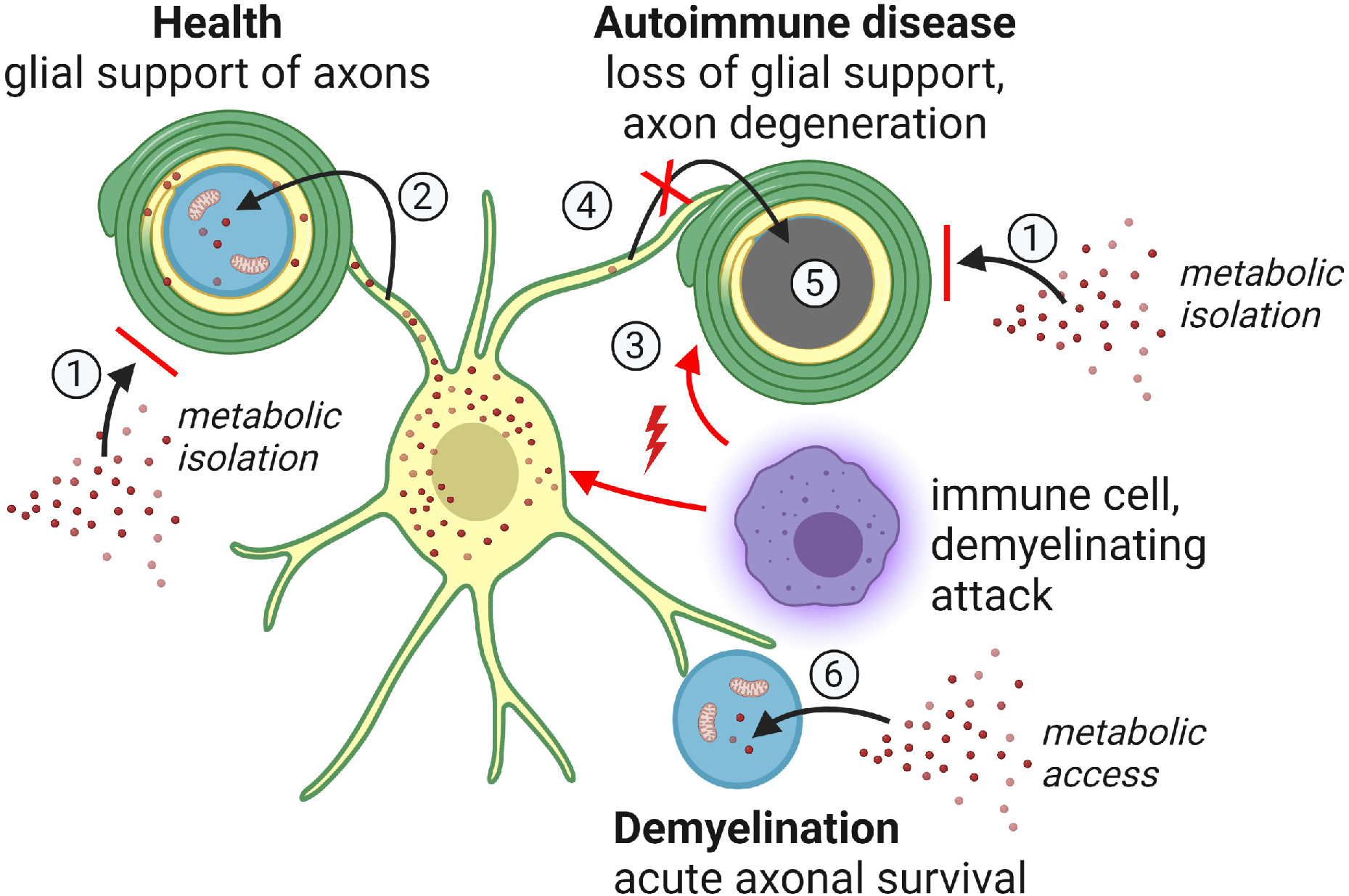
Working model on the role of myelin in autoimmune disease. Myelinated axons are metabolically isolated from the environment by the myelin sheath (1), and are therefore dependent on metabolic support by oligodendrocytes (2). An autoimmune attack against myelinating oligodendrocytes by immune cells (3) perturbs glial support (4). Axons that are hence trapped in metabolic isolation are prone to degenerate (5). Rapid demyelination, in turn, facilitates metabolic access of axons to the extracellular environment, which acutely secures axonal survival (6).

## Notes

### Competing Interest Statement

The authors have declared no competing interest.

